# Seasonal zonation patterns of the sandy beach bivalve *Donax deltoides* (Bivalvia: Donacidae) in subtropical eastern Australia

**DOI:** 10.1101/610576

**Authors:** Stephen Totterman

## Abstract

Seasonal low tide zonation patterns of the sandy beach bivalve *Donax deltoides* are described from eleven beaches in subtropical eastern Australia. Ten low tide transects were sampled at equal along shore intervals along the length of each beach. Three quadrats were sampled at each of seven across shore levels on each transect, from the high tide drift line, across the intertidal zone and to the bottom of the low tide swash zone. Zone counts showed that *D. deltoides* shifted towards the low tide swash zone in summer and buried in the intertidal zone in winter. It is proposed that *D. deltoides* moves down shore in summer to avoid thermal stress in the intertidal zone. This seasonal pattern is important when planning surveys for this species.

## 1. Introduction

This paper concerns the across shore distribution or “zonation” of *Donax deltoides* at low tide. Commonly known as the “pipi”, *D. deltoides* is a large sandy beach bivalve or “surf clam” found along the eastern and southern coasts of Australia, from Fraser Island, Queensland, across New South Wales (NSW) and Victoria, to the Murray River, in South Australia. *D. deltoides* grows to a maximum length of 75 mm and can develop large populations on some beaches, where they are harvested by commercial and recreational fishers (Ferguson *et al.* 2014). Owing to its large size, *D. deltoides* can make up a large component of the macrobenthic biomass over a wide range of densities (*e.g.* McLachlan *et al.* 1996) and has an important role in the ecology of sandy beaches. Efforts to quantify *D. deltoides* populations have recently increased following growth in commercial and recreational fisheries for this species (Ferguson *et al.* 2015).

Two key adaptations of “surf clams”, a functional group comprising numerous *Donax* species and also including some species from the genera *Mesodesma, Paphies, Donacilla* and others, for the beach environment are the muscular foot for rapid burrowing and “swash riding” behaviour for moving up and down the beach face (Ellers 1995; McLachlan *et al.* 1995). Tidal migrations are common (Ansell and Trueman 1973), where large numbers of clams remain in the swash zone for all or part of the tidal cycle. Potential benefits of tidal migration for surf clams include increased feeding time in the swash zone, where phytoplankton and organic detritus are held in suspension by wave action, and reduced predation risk from marine predators. Costs include energy expenditure for repeated emergence and burial when migrating, increased predation risk from swash zone predators and risk of stranding on ebb tides (Gibson 2003). Movements and zonation of surf clams can be further influenced by other physical and biological factors that can affect burrowing and mobility (McLachlan and Young 1982; Soares *et al.* 1996; De la Huz *et al.* 2002) or increase competition for space or food resources (Ansell 1983; Dugan *et al.* 2004). Varying conditions result in surf clam aggregations that vary across small and large spatial and temporal scales (Donn 1990; Defeo and McLachlan 2005) including seasonal and other rhythmic patterns (Donn *et al.* 1986; Defeo *et al.* 1986).

Spatial and temporal variation in aggregations is an important sampling issue for surf clams. Clams are commonly sampled at low tide by excavating and sieving quadrats on across shore transects. A common defect in surf clam sampling designs is to neglect along shore variation and sample only one transect, which may not necessarily be representative of the population of clams on the beach (Murray-Jones 1999). Murray-Jones (1999) previously attempted to study zonation of *D. deltoides* on east Australian beaches using single transects but those results were highly variable and did not show clear patterns. This paper presents the first study of low tide zonation of *D. deltoides* with transect sampling across the intertidal and low tide swash zones, replicate transects per beach, replicate beaches and covering all seasons.

## 2. Material and Methods

### 2.1 Study sites

Intermediate energy, “microtidal”, wave-dominated beaches are common along the coast of NSW, with double-bar systems on longer beaches (Short 2007). East (41%) to south-easterly swells (40%) and moderate waves (mean height 1.6 m, mean period 10 s) predominate. Tides are semi-diurnal and unequal (mean range 1.3 m, maximum range 2 m at Port Jackson with < 0.2 m range difference along the coast).

Eleven beaches were sampled in northern NSW from the Tweed River to the Wooli River (Fig. 1). Beaches were delimited by natural breaks such as headlands, rivers and rocky intervals. Beaches where *D. deltoides* was known to be abundant were preferred, including those identified in a previous regional study by Owner and Rohweder (2003). South Ballina, which had the highest and increasing *D. deltoides* counts was sampled four times (Table 1). Surveys were performed in all months except for March and May.

**Table 1.** Beach summaries, ordered north to south (Fig. 1). Each beach was sampled with 10 transects in the months indicated (South Ballina was sampled on four occasions). Latitude is given for the southern end of each beach. Beach morphodynamics from Short (2007) are, in order of increasing energy: LTT = low tide terrace, TBR = transverse bar and rip, RBB = rhythmic bar and beach. Medium sand has a grain size 0.25–0.5 mm on the Wentworth scale.

| Beach | Sampled | Latitude (S) | Inner/outer bars | Length (km) | Mean width (m) | Mean aspect (°) | Mode grain size | Mean slope (cm/10 m) | Mean water (°C) |
| --- | --- | --- | --- | --- | --- | --- | --- | --- | --- |
| Letitia Spit | Apr-15 | 28°11'47" | TBR/RBB | 2.8 | 63 | 68 | Medium | 41 | 24 |
| North | Jan-16 | 28°21'28" | TBR/RBB | 2.3 | 79 | 93 | Medium | 50 | 25 |
| Wooyung | Dec-13 | 28°32'10" | TBR/RBB | 16.6 | 77 | 97 | Medium | 37 | 21 |
| Seven Mile | Jul-14 | 28°47'06" | TBR/RBB | 6.7 | 92 | 109 | Medium | 46 | 20 |
| South Ballina | Aug-13 | 29°01'24" | TBR/RBB | 19.1 | 61 | 126 | Medium | 42 | 20 |
| South Ballina | Jan-14 | 29°01'24" | TBR/RBB | 19.1 | 75 | 125 | Medium | 51 | 24 |
| South Ballina | Sep-14 | 29°01'24" | TBR/RBB | 19.1 | 61 | 129 | Medium | 36 | 20 |
| South Ballina | Feb-15 | 29°01'24" | TBR/RBB | 19.1 | 68 | 125 | Medium | 44 | 27 |
| Evans Head | Aug-15 | 29°06'08" | TBR/RBB | 4.6 | 98 | 103 | Medium | 29 | 20 |
| Ten Mile | Apr-14 | 29°21'34" | TBR/RBB | 12.5 | 80 | 96 | Medium | 30 | 22 |
| Iluka | Dec-14 | 29°25'16" | TBR | 2.7 | 79 | 108 | Medium | 53 | 21 |
| Sandon | Oct-13 | 29°40'24" | LTT-TBR/RBB | 6.3 | 62 | 99 | Medium | 48 | 21 |
| Illaroo | Nov-14 | 29°45'32" | TBR/RBB | 8.8 | 76 | 122 | Medium | 47 | 22 |
| Wooli | Jun-14 | 29°53'13" | LTT-TBR/RBB | 6.7 | 47 | 109 | Medium | 85 | 21 |

**Fig. 1.**
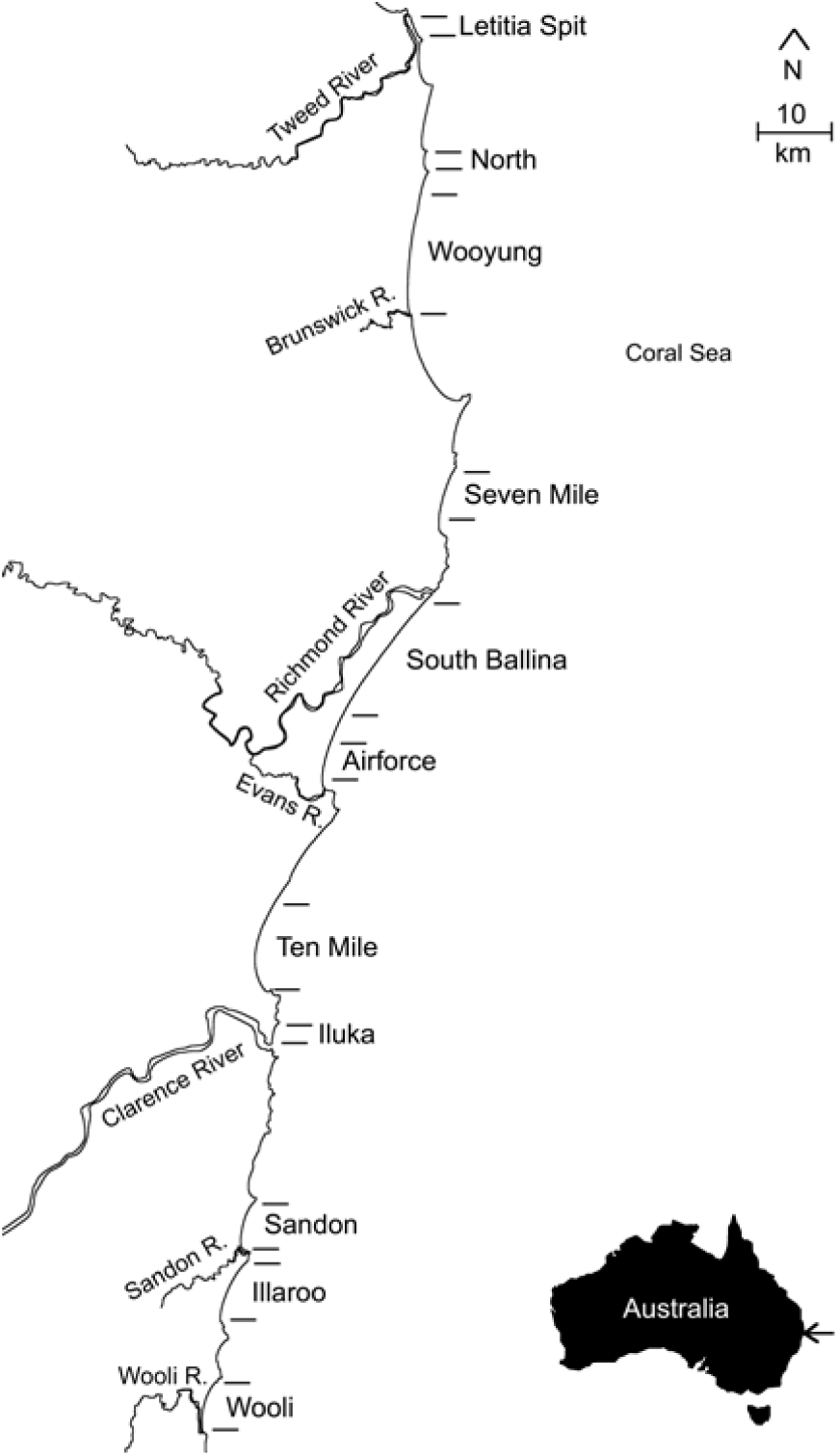
Location of 11 study beaches. The inset map shows the study region on the east coast of Australia (arrowed).

This study applied a tide-based, four-zone scheme to describe low tide zonation (Fig. 2). The “supertidal zone” extends down to the high tide “drift”, “debris” or “wrack” line, defined by the maximum run-up of waves at high tide. The “intertidal zone” extends from the high tide drift line down to the top of the “low tide swash zone”. The sand here is moist and can be compact and hard on beaches with fine–medium grain sand (< 0.5 mm). The “swash zone” is repeatedly flooded and exposed by the swash and backwash. The sand here is water saturated and thixotropic. Surf clams are swash riders, swash zone feeders and burrow most rapidly in thixotropic sand. Thus, the top of the swash zone is a biologically reasonable zone boundary (more so than the “effluent line”, which is observed where water seeps out of the intertidal zone). The swash zone moves up and down the beach face with the tidal range. This study sampled the swash zone only at low tide. Below the low tide swash zone is the “subtidal zone”, which is continuously covered by water.

**Fig. 2.**
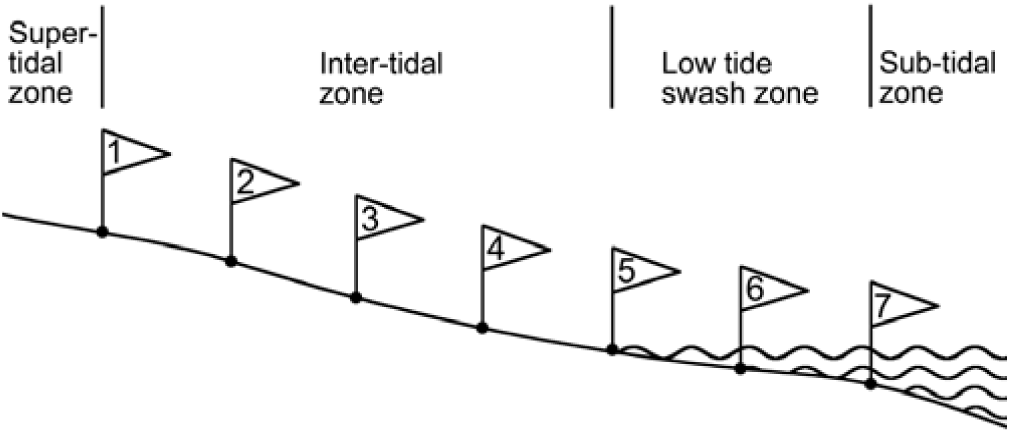
Beach profile illustrating low tide zonation scheme used in this study and transect sampling levels. Levels 1–4 are in the intertidal zone. Levels 5–7 are in the low tide swash zone.

### 2.2 Transect sampling

Each beach was sampled with ten transects, equally spaced along the length of the beach starting at a random offset from the southern end (distance 0 km). Sampling occurred over several days because no more than two transects could be sampled during one low tide period. No particular attention was given towards spring-neap or semi-diurnal variation in the tidal range. It was assumed that *D. deltoides* undergo tidal migrations to optimise their across shore location relative to varying beach conditions.

A fixed area transect design with variable across shore level intervals was used to consistently position transect levels relative to the limits of the intertidal and swash zones, taking into account variable beach width at transect locations, varying tidal range and varying swash run up (Fig. 2). Transects extended from the most recent high tide drift line to the bottom of the low tide swash zone (at the average limit of the backwash). Level one was at the drift line, levels 2–4 were equally spaced in the intertidal zone, level five was at the top of the low tide swash zone (at the average limit of swash run up), level six was in the middle of the swash zone and level seven was at the bottom of the swash zone (average limit of backwash) (Fig. 2). Three replicate quadrats were sampled in a direction parallel to the shoreline at each of levels 1–6, with five paces between quadrats. Excavation of large quadrats underwater is impossible and three “feet digging” plots were sampled in place of quadrats at level seven (see below).

The mean transect level interval was 9 m (range 3–23 m). Schlacher *et al.* (2008), recommended 7–27 transect levels (increasing with beach width) for sampling sandy beach macrofauna. Totterman (2019), in the context of abundance estimation, demonstrated that the level interval should be no greater than 0.5× the width of the macrofauna band. Seven transect levels was adequate for abundance estimation except that undersampling can possibly occur if macrofauna bands are narrow and the beach is wide. Concerning zonation studies, it is possible that a macrofauna band that is narrower than the transect level interval is completely missed. This concern is less critical for beach scale sampling however, because the chance of repeatedly failing to intersect aggregations with several transects is low. Moreover, this study examines relative counts in the low tide intertidal and swash zones (see 2.4 Statistical analysis) and transect counts near zero do not bias the beach (population) scale across shore distribution pattern.

A square, sheet metal, 0.1 m^2^ quadrat (sides 31.6 cm long and 10 cm deep) was used (James and Fairweather 1995). Sand was excavated first to 10 cm and then separately from 10–20 cm. A hand shovel was used for gauging the 10–20 cm depth. Excavated material was washed through 6 × 6 millimetre plastic sieves using buckets of seawater except that “finger raking” (James and Fairweather 1995) was used to rapidly inspect the 10–20 cm depth in saturated sand, where the sides of the quadrat hole were prone to collapsing. Finger raking of flooded quadrats involves tactile detection and dislodgment of buried clams, which tend to float to the surface. Finger raking was also frequently used for the 0–10 cm depth at level six in the swash zone, where waves disrupts sampling efforts. Clams were measured with callipers to the nearest millimetre (maximum shell length) and then released. Clams broken at the edges of the quadrat and not measurable or any clams washed away in the swash zone and lost were added to the count of those measured. Clams smaller than ten millimetres passed through the diagonal apertures of the sieve mesh and were recorded only as present/absent.

Feet digging or “feet twisting” is commonly used by recreational fishers collecting *D. deltoides* in the swash zone (see Jaramillo *et al.* 1994 for a previous application of this method). Twisting one’s legs and feet from the hips down while pivoting on the balls of the feet, causes the feet to dig into the sand. Erosion of sand from around the feet by swash and backwash speeds the process. Clams disturbed in the vicinity of the feet rise to the surface where they can be picked up (similar to the “finger raking” method of James and Fairweather 1995). Large and more firmly anchored clams can be felt with the feet and recovered by hand. The area sampled by one feet digging plot is approximately 0.1 m^2^ and the depth is approximately 10 cm.

### 2.3 Physical variables

Physical variables thought to influence zonation for *D. deltoides* were temperature, beach slope and sand grain size. High temperatures are lethal for surf clams (Ansell *et al.* 1980; Ansell and McLachlan 1980) and low temperatures slow burrowing rates (McLachlan and Young 1982; Donn 1990). Steep slopes can hinder movements up shore and promote movements down shore. Coarse sand >0.5 mm slows burrowing rates for larger clams (De la Huz *et al.* 2002). Biological variables such as stranded kelp (Soares *et al.* 1996) and interspecific competition (Ansell 1983; Dugan *et al.* 2004) did not feature on those beaches sampled in this study.

Beach lengths were measured and transects were located by GPS. Physical variables measured at each transect were: beach width (from the base of the incipient fore dune to the middle of the low tide swash zone, using a marked rope), beach aspect (using GPS and walking down the transect) and air temperature. Variables measured at each transect level were: sand temperature at 10.5 cm depth, water temperature (at level 7), slope (height difference) between graded poles ten metres apart (Emery 1961) and sand grain size (a handful of sand was scooped from the surface and classified on the Wentworth scale by visual comparison to graded samples on a sand gauge). Grain size assessments ignored any shell fragments present. Slope could not be measured at level seven because of water depth and waves. The same digital thermometer probe was used for measuring sand, water and air temperatures.

### 2.4 Statistical analysis

The transect sampling design used gives a spatial hierarchy of counts: 1) counts of *D. deltoides* from 0–10 cm or 10–20 cm within each quadrat, 2) quadrat counts (from 0–20 cm), 3) level counts (the sum of the three quadrat counts within a transect level), 4) transect counts (the sum of all levels within a transect), and 5) beach counts (the sum of all transects within a beach). A detailed hierarchical analysis (beach/transect/level/quadrat/depth) was not possible because of low counts at several beaches and many zero counts at the three smallest scales. Rather, summation of counts at larger scales of investigation eliminates pseudoreplication (Millar and Anderson 2004), reduces variance and simplifies analysis and reporting of results (Murtaugh 2007). Level counts were grouped into intertidal (the sum of levels one to four) and low tide swash (the sum of levels five to seven) zones (Fig. 1). As there were zero *D. deltoides* at level one, the sampling effort was effectively balanced at three levels in each zone. To emphasise beach (population) scale zonation and further reduce variance, zone counts were summed for all ten transects within each beach. Thus, beach scale zone counts indicate the relative abundance of *D. deltoides* in the intertidal and swash zones for a specific population (beach) during a particular sampling period (within a month).

Like counts, physical variables were summarised at the mid-intertidal (level three) and mid-swash (level six) zones at the beach scale. Physical variables measured at adjacent levels were strongly correlated (*r* ≥ 0.94 for temperature, *r* ≥ 0.92 for slopes) and results were not sensitive to selecting mid-intertidal and mid-swash values as representative of those two zones. Summaries of physical variables were based on only those transects with *D. deltoides* present as zero counts do not contribute to zonation patterns. Level temperatures within beaches were summarised as maximums rather than means because of diurnal variation in temperature. Transects were often sampled early in the morning when sand temperatures are lower than near midday. Moreover, maximum temperature is biologically reasonable (*D. deltoides* could be sensitive to maximum intertidal temperatures rather than means) and practical (maximum temperatures are more clearly defined than are means). Level slopes within beaches were summarised as means. Level grain sizes within beaches were summarise as modes (the Wentworth scale is logarithmic and means are not appropriate).

Relationships between zonation in *D. deltoides* counts and physical variables were examined using logistic regression models with quasibinomial errors to account for overdispersion. The response variable is the proportion of counts in the low tide swash zone (*i.e.* swash count / (swash + intertidal count)). The logit link function linearises the response and weighted regression is performed using the individual sample sizes (*i.e.* swash + intertidal counts) as weights (Crawley 2007). Predictor variables are maximum temperature, mean slope and mode grain size for the mid-intertidal and mid-swash zones. Regression models were compared using information theoretic methods adjusted for overdispersion using quasi-likelihood theory and corrected for small sample sizes (Anderson *et al*. 1994). Smaller overdispersion (variance inflation factor, VIF < 5–10) suggests an adequate model structure and smaller corrected quasi Akaike’s Information Criterion (QAIC_c_) indicates a more parsimonious model. Goodness-of-fit is summarised as deviance pseudo-*R*^2^: *R*^2^_*D*_ = *1 –* ln(*L*_*β*_) / ln(*L*_*0*_), where *L*_*β*_ is the likelihood of the data given the fitted model of interest and *L*_*0*_ is the likelihood of the data given the null (intercept-only) model (McFadden 1974). All statistical analyses were performed using R version 3.3.2 (R Core Team 2016). Logistic regressions were computed using the glm function in the stats package version 3.3.2. Median zonation thresholds (where 50% of clams are in the swash zone) were computed using the dose.p function in the MASS package version 7.3-45 (Venables and Ripley 2002).

## 3. Results

Abundance of *D. deltoides* was low on most beaches and most clams sampled were adults. Total beach counts ranged from three clams at Iluka to 184 at South Ballina and the median count was 31. Median shell lengths were > 37 mm, where 50% of *D. deltoides* are sexually mature (Murray-Jones 1999), on all beaches except Wooli, where the median was 31 mm.

Transect quadrats recovered 669 of 677 *D. deltoides* (99%) at 0–10 cm depth. *D. deltoides* was found in the low tide swash and intertidal zones but not at the high tide drift line (level one). Unimodal across shore count distributions within beaches indicate that *D. deltoides* populations maintained some optimum position relative to the tidal range (Fig. 3). Variable level counts between transects, within beaches indicate alongshore variation in abundance (*i.e.* variable counts at the transect scale; boxes and whiskers in Fig. 3). Level counts showed seasonal changes in low tide zonation of *D. deltoides*. Higher counts occurred in the low tide swash zone (levels 5–7) and apparently into the subtidal in the summer months of January and February (Figs. 3a1, b1). Higher counts occurred in the intertidal zone (levels 2–4) in the winter and early spring months of June–August (Figs. 3c1, d1, e1). Zonation of new recruits < 10 mm was similar to counts of larger *D. deltoides* except that these tiny clams can be overlooked in the swash and subtidal zones when sampling with finger raking or feet digging (James and Fairweather 1995).

**Fig. 3.**
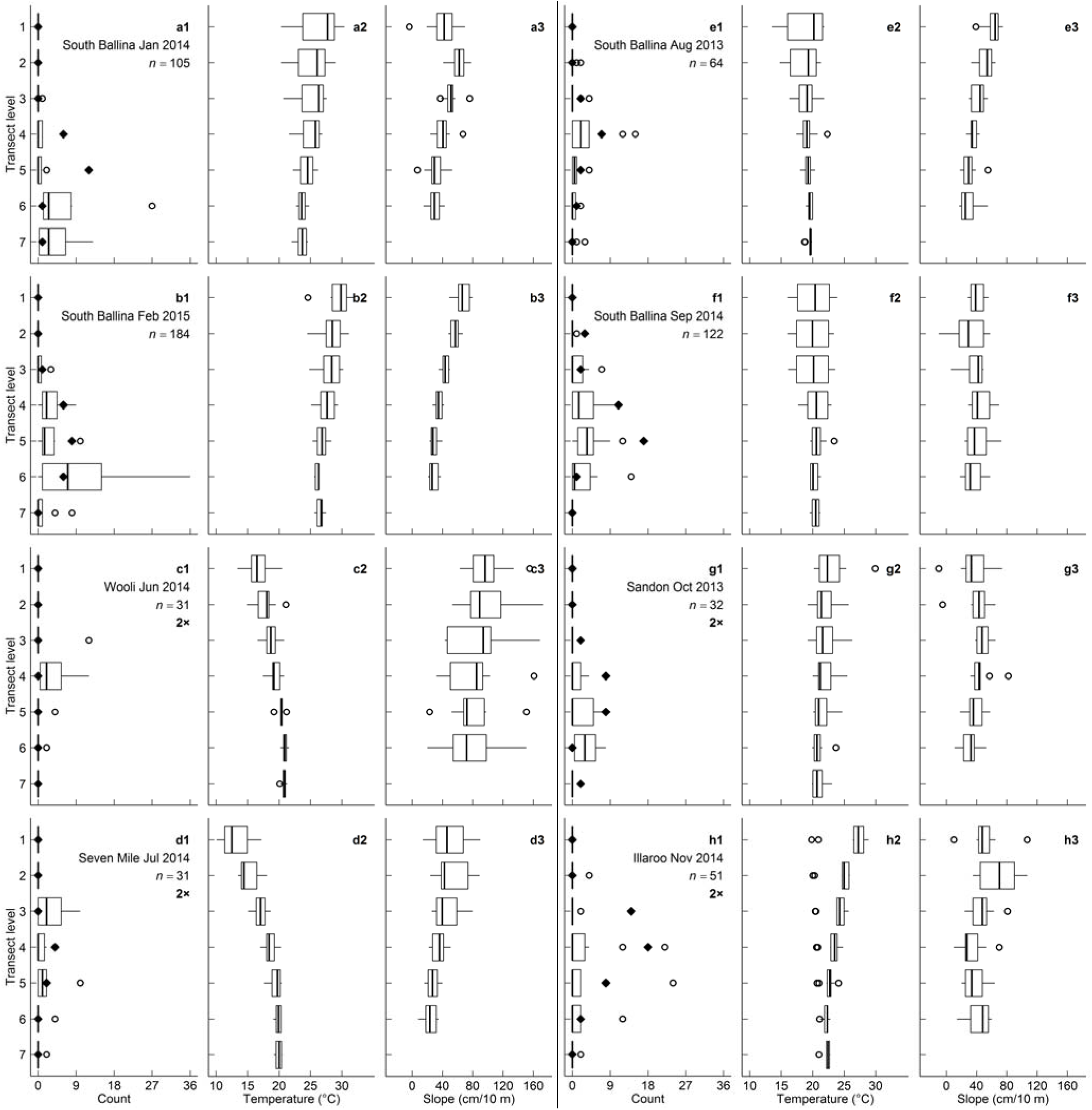
Box and whisker plots of transect level *D. deltoides* counts, temperatures and slopes for beaches with *n* > 30 *D. deltoides*. Beaches with lower counts are not presented because clam aggregations were more weakly expressed at the transect scale (see Fig. 4 for complete beach-scale results). Plots are grouped by beaches across rows (1–3 and 4–6) and columns are ordered by month (a–h). Beaches with smaller counts are presented with 2× expanded scales. Level one was at the high tide drift line, level five was at the top of the low tide swash zone and level seven was at the bottom of the swash zone (Fig. 2). Level seven slopes were not measured because of water depth and waves. Boxes show the first quartile, median and third quartile, whiskers extend to a maximum of 1.5 times the interquartile range and data outside the whiskers are plotted as individual points (white-filled circles). Frequencies of quadrats with *D. deltoides* < 10 mm present are plotted as diamonds (zero were observed at Wooli).

Median level temperatures showed opposite gradients in summer/winter. Temperature decreased towards the low tide swash zone in October–February (Figs. 3a2, b2, h2) and increased towards the swash zone in June and July (Figs. 3c2, d2). The high tide drift line (level one) had the widest temperature range, where the minimum recorded was 10.1°C and the maximum was 33.4°C. Temperature variability decreased towards the swash zone and subtidal. The lowest recorded water temperature (level seven) was 18.4°C and the highest recorded water temperature was 27.5°C.

Variable profiles between beaches largely resulted from varying physical characteristics (Table 1) and no seasonal variation was apparent (Fig. 3). Wooli had a substantially steeper beach slope than did other beaches. Grain size increased down shore, although this pattern was not clearly resolved on the Wentworth scale. The modal grain size was medium (0.25–0.5 mm) across all beaches and all transect levels except Wooli, which had very coarse sand (1–2 mm) at level six and very fine gravel (2–4 mm) at level seven and Letitia Spit, which had fine sand (0.125–0.25 mm) at level one.

Four logistic regression models for zonation in *D. deltoides* swash and intertidal counts were examined: 1) mid-intertidal maximum temperature (level three), 2) mid-swash maximum temperature (level six), 3) mid-intertidal mean slope, and 4) mid-swash mean slope. Sand grain size, for which there was almost no measured variation, was excluded. The mid-intertidal temperature model was selected for having the smallest overdispersion (VIF = 5.5) and largest goodness-of-fit statistic (*R*^*2*^_*D*_ = 0.52). Quasi-AIC for the mid-swash temperature model was smaller than that for the mid-intertidal temperature model (ΔQAIC_c_ = 2.3), however overdispersion was larger (VIF = 7.2) which resulted in a large adjustment to AIC for that model. Mid-intertidal temperature and slope models are compared in Fig. 4. Similar temperature (seasonal) zonation patterns across the wide range of beach scale counts indicate that zonation for *D. deltoides* was not sensitive to abundance.

**Fig. 4.**
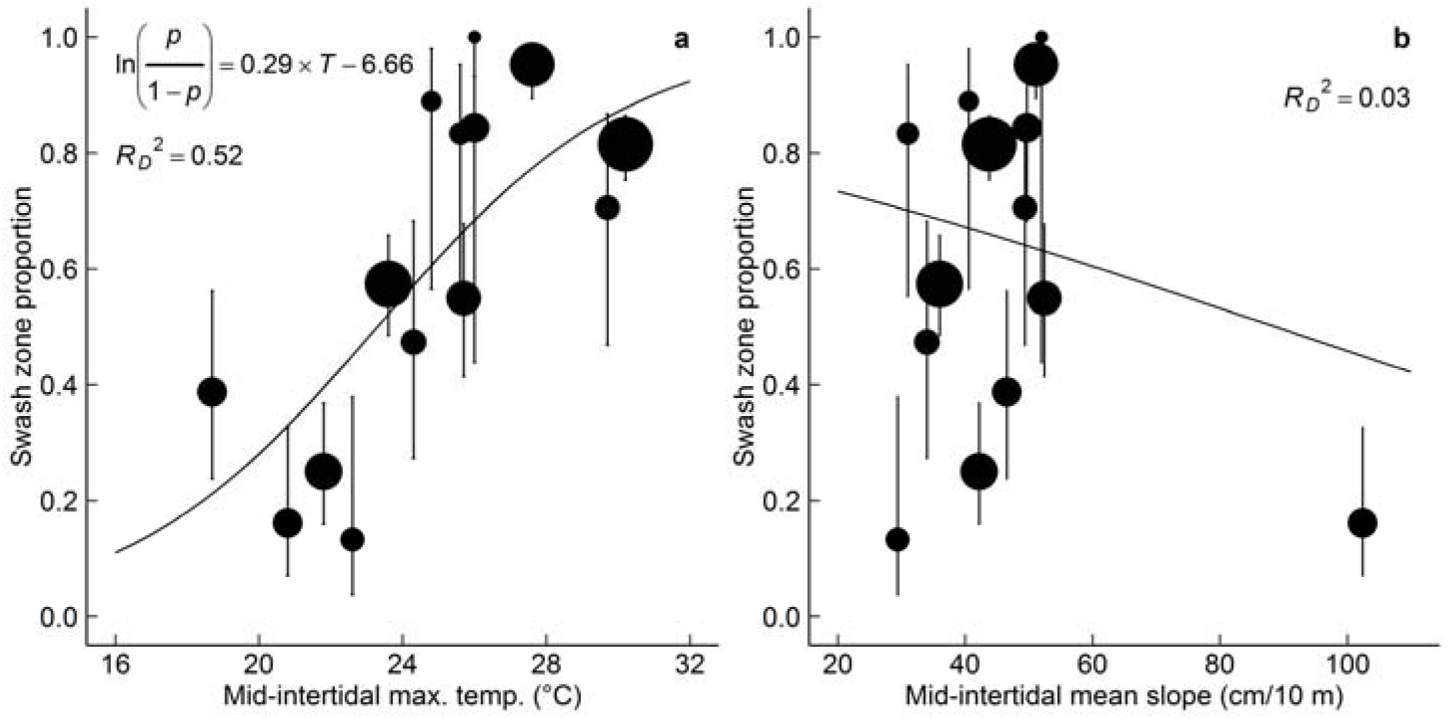
Mid-intertidal (level three) temperature (a) and slope (b) logistic regression models (curved lines) for *D. deltoides* zonation (proportion of beach scale total counts in the swash zone). Points are 14 beaches (diameters are scaled to total counts). Vertical lines are 95% Wilson confidence intervals (Brown *et al.* 2001).

## 4. Discussion

Sampling of 11 beaches with 10 replicate transects per beach showed that counts of *D. deltoides* varied at multiple spatial and temporal scales: between beaches, along shore between transects within beaches, across shore between zones within beaches and across shore between seasons within zones. Seasonal zonation patterns have not previously been described for *D. deltoides*. One reason could be inadequate replication of transects within beaches (Murray-Jones 1999). Another is that zonation is likely to differ in other regions, like the cold/warm water contrast that has been reported for *D. serra* populations in South Africa (Donn 1990). Further studies could investigate zonation of *D. deltoides* on cold water beaches on the southern coast of Australia.

Results from this study can improve quadrat-based sampling designs for *D. deltoides*. Only 1% of clams were recovered from 10–20 cm, the same as 1% from 10–20 cm in James and Fairweather (1995) and zero from > 15 cm in Murray-Jones (1999). Quadrats deeper than 10 cm are unnecessary. Clams were absent at the high tide drift line, the same as James and Fairweather (1995; 1996) and Murray-Jones (1999). The high tide drift line and certainly the supertidal zone does not have to be sampled. Clams typically buried in the intertidal zone during winter and early spring, where quadrat sampling is more easily performed than in the swash zone during summer.

Designs that sample only the intertidal or only the low tide swash zone can produce biased results. Gray (2016a, 2016b) studied the effects of commercial fishing on *D. deltoides* on six NSW beaches including South Ballina, Ten Mile, Sandon and Illaroo that were sampled in this study. Beaches were sampled at low tide in April/May (autumn), July/August (winter) and in October/November (early summer) by two different methods: 1) 30-second “hand digging” plots in the swash zone (Gray *et al.* 2014), and 2) 0.1 m^2^ quadrats in the intertidal zone. Mean counts from these two zones showed contrasting seasonal patterns. Intertidal zone counts peaked in April/May or July/August and low tide swash zone counts peaked in October/November. No consistent harvest effect was detected. It could be that Gray’s (2016a, 2016b) counts were biased by seasonal changes in low tide zonation of *D. deltoides* like that observed in this study.

Seasonal zonation patterns in *D. deltoides* were associated with seasonality in temperature. It is proposed that *D. deltoides* avoids high temperatures in the intertidal zone by moving down shore in summer. Thermal tolerances for *D. deltoides* are unknown. Upper median lethal temperatures for five other *Donax* species range from 25–33°C at 24 hours exposure (Ansell and McLachlan 1980; Ansell *et al.* 1980). Bivalves begin to experience heat stress at sub-lethal temperatures (Brown *et al.* 1989) and logistic regression models from this study predict that *D. deltoides* shifts towards the swash zone when maximum intertidal temperatures exceed 23°C. Burrowing more deeply can insulate surf clams from temperature extremes (Brown *et al.* 1989), however this has not been observed for *D. deltoides*. Rather, *D. deltoides* buried in the intertidal zone at low tide sometimes rise upwards, pushing the sand above them to form sand “caps” on the surface. This vertical movement could be a response to thermal stress, like that observed for some other *Donax* species (Ansell *et al.* 1980; Brown *et al.* 1989) rather than vehicle traffic (Murray-Jones 1999) because sand caps can also occur where vehicles have not passed (pers. obs.). Further studies are required to test whether seasonal zonation patterns for *D. deltoides* actually are responses to changes in intertidal temperatures. Laboratory measurement of thermal tolerances for *D. deltoides* would be useful.

*D. deltoides* typically buried in the intertidal zone at low tide in winter and early spring rather than stay in swash zone. Water temperatures were > 15°C, where Donn (1990) reported slowed burrowing rates and *D. serra* aggregations in the swash zone. One cost of staying in the intertidal zone is reduced feeding time due to a shorter period of inundation. Two benefits could be lower energy costs compared to tidal migration and lower predation risk from Australian pied oystercatcher *Haematopus longirostris* birds which forage for *D. deltoides* in the swash zone (Owner and Rohweder 2003). Ansell and Trueman (1973) estimated for two *Donax* species on tropical beaches that energy expenditure for maintaining position by repeated burrowing in response to erosion and deposition of sand by waves was greater than that for tidal migration. However, McLachlan and Hanekom (1979) suggested that the opposite could be true for large clams on wider, gently sloping, more dissipative beaches, for which migratory movements are longer and where the swash environment is relatively calm.

Observed seasonal zonation patterns for *D. deltoides* agree with James and Fairweather’s (1995; 1996) results from Catherine Hill Bay (33° 09’ S), on the NSW central coast. In February 1994 (summer), they sampled ten low tide transects and found 80% of *D. deltoides* individuals in the swash zone. In May (early winter), they sampled two transects and found clams aggregated in the mid-intertidal. In June (winter), they sampled six transects and again found clams in the mid-intertidal. In October (late spring), they sampled two transects and found clams aggregated in the low-intertidal, further down shore than in June. Observed seasonal zonation patterns also agree with the behaviour of commercial fishers in NSW, who typically harvest *D. deltoides* from the intertidal zone at low tide, searching for patches of sand caps that can form above aggregations of *D. deltoides*. Murray-Jones and Steffe (2000) observed that commercial fishing effort and catch on Stockton Beach, NSW were highest in winter and that clams had become scarce in summer. While fishing could have substantially decreased abundance, clams could also have moved into the low tide swash zone and become more difficult to find and harvest in summer (“cockle rakes” that are used in the swash zone on beaches in South Australia are prohibited in NSW; Ferguson *et al.* 2014).

Seasonal zonation patterns for *D. deltoides* described in this study are important when planning surveys for this species on eastern Australian beaches. However, these patterns concern “average”, beach scale zonation. Varying conditions at smaller spatial and temporal scales (Defeo and McLachlan 2005) mean that zonation at the transect sampling unit is less predictable. Sampling across the intertidal and swash zones is always recommended to prevent bias.

## Abbreviations

NSW: New South Wales
QAIC: quasi Akaike’s information criterion
VIF: variance inflation factor

## Acknowledgements

Catch, measure and release sampling of *D. deltoides* was permitted under Section 37 of the Fisheries Management Act 1994, permit number P13/0024-1.0 issued by Fisheries NSW, Department of Primary Industries, NSW Government.

This paper was submitted for publication twice resulting in feedback from three reviewers. While one reviewer recognised the practical importance of this work, comments from the other two, and even from Journal editors, showed a rather arrogant and elitist attitude. The field of sandy beach ecology is specialised and it seems the relevant journals largely serve an elite group of researchers.

## References

Anderson, D.R., Burnham, K.P., White, G.C., 1994. AIC model selection in overdispersed capture-recapture data. Ecology 75: 1780–1793. https://doi.org/10.2307/1939637.

Ansell, A.D., 1983. The biology of the genus *Donax*, in: McLachlan, A., Erasmus, T. (Eds.), Sandy Beaches as Ecosystems, Developments in Hydrobiology, vol. 19. Springer, Dordrecht, pp. 607–635. https://doi.org/10.1007/978-94-017-2938-3_46.

Ansell, A.D., Trueman, E.R., 1973. The energy cost of migration of the bivalve *Donax* on tropical sandy beaches. Marine Behaviour and Physiology 2: 21–32. https://doi.org/10.1080/10236247309386914.

Ansell, A.D., Barnett, P.R.O., Bodoy, A., Massé, H. 1980. Upper temperature tolerances of some European molluscs. II. *Donax vittatus, D. semistriatus* and *D. trunculus*. Marine Biology 58: 41–46. https://doi.org/10.1007/BF00386878.

Ansell, A.D., McLachlan, A., 1980. Upper temperature tolerances of three molluscs from South African sandy beaches. Journal of Experimental Marine Biology and Ecology 48: 243–251. https://doi.org/10.1016/0022-0981(80)90079-9.

Brown, A.C., Stenton-Dozey, J.M.E., Trueman, E.R., 1989. Sandy-beach bivalves and gastropods: a comparison between *Donax serra* and *Bullia digitalis*, in: Blaxter, J.H.S., Southward, A.J. (Eds.), Advances in Marine Biology, vol. 25. Academic Press, London, pp. 179–247. https://doi.org/10.1016/S0065-2881(08)60190-X.

Brown, L.D., Cai, T.T., DasGupta, A., 2001. Interval estimation for a binomial proportion. Statistical Science 16: 101–133. http://www.jstor.org/stable/2676784.

Crawley, M.J., 2007. The R Book. John Wiley and Sons Ltd, Chichester.

Defeo, O., Layerle, C., Masella, A., 1986. Spatial and temporal structure of the yellow clam *Mesodesma mactroides* (Deshayes, 1854) in Uruguay. Medio Ambiente 8: 48–57.

Defeo, O., McLachlan, A., 2005. Patterns, processes and regulatory mechanisms in sandy beach macrofauna: a multi-scale analysis. Marine Ecology Progress Series 295: 1–20. https://doi.org/doi:10.3354/meps295001.

De la Huz R., Lastra, M., López, J., 2002. The influence of sediment grain size on burrowing, growth and metabolism of *Donax trunculus* L. (Bivalvia: Donacidae). Journal of Sea Research 47: 85–95. https://doi.org/10.1016/S1385-1101(02)00108-9.

Donn, T.E., Clarke, D.J., McLachlan, A., DuToit, P., 1986. Distribution and abundance of Donax serra Röding (Bivalvia: Donacidae) as related to beach morphology. I. Semilunar migrations. Journal of Experimental Marine Biology and Ecology 102: 121–131. https://doi.org/10.1016/0022-0981(86)90171-1.

Donn, T.E., 1990. Zonation patterns of *Donax serra* Röding (Bivalvia: Donacidae) in southern Africa. Journal of Coastal Research 6: 903–911. http://www.jstor.org/stable/4297763.

Dugan, J.E., Jaramillo, E., Hubbard, D.M., Contreras, H., Duarte, C., 2004. Competitive interactions in macroinfaunal animals of exposed sandy beaches. Oecologia 139: 630–640. https://doi.org/10.1007/s00442-004-1547-x.

Ellers, O., 1995. Behavioral control of swash-riding in the clam *Donax variabilis*. The Biological Bulletin 189: 120–127. https://doi.org/10.2307/1542462

Emery, K.O., 1961. A simple method of measuring beach profiles. Limnology and Oceanography 6: 90–93. https://doi.org/10.4319/lo.1961.6.1.0090.

Ferguson, G., Johnson, D., Andrews, J., 2014. Pipi *Donax deltoides*, in Flood, M., Stobutzki, I.,Andrews, J., Ashby, C., Begg, G., Fletcher, R., Gardner, C., Georgeson, L., Hansen, S., Hartmann, K., Hone, P., Horvat, P., Maloney, L., McDonald, B., Moore, A., Roelofs, A., Sainsbury, K., Saunders, T., Smith, T., Stewardson, C., Stewart, J., Wise, B. (Eds.), Status of Key Australian Fish Stocks Reports 2014. Fisheries Research and Development Corporation, Canberra.

Ferguson, G.J., Ward, T.M., Gorman, D., 2015. Recovery of a surf clam *Donax deltoides* population in Southern Australia: successful outcomes of fishery-independent surveys. North American Journal of Fisheries Management 35: 1185–1195. https://doi.org/10.1080/02755947.2015.1091408.

Gibson, R.N., 2003. Go with the flow: tidal migration in marine animals, in: Jones, M.B., Ingólfsson, A., Ólafsson, E., Helgason, G.V., Gunnarsson, K., Svavarsson, J. (Eds), Migrations and Dispersal of Marine Organisms. Developments in Hydrobiology, vol. 174. Springer, Dordrecht, pp. 153–161. https://doi.org/10.1007/978-94-017-2276-6_17.

Gray, C.A., Johnson, D.D., Reynolds, D., Rotherham, D., 2014. Development of rapid sampling procedures for an exploited bivalve in the swash zone on exposed ocean beaches. Fisheries Research 154: 205–212. https://doi.org/10.1016/j.fishres.2014.02.027.

Gray, C.A., 2016a. Assessment of spatial fishing closures on beach clams. Global Ecology and Conservation 5: 108–117. https://doi.org/10.1016/j.gecco.2015.12.002.

Gray, C.A., 2016b. Effects of fishing and fishing closures on beach clams: experimental evaluation across commercially fished and non-fished beaches before and during harvesting. PLoS ONE 11: e0146122. https://doi.org/10.1371/journal.pone.0146122.

James, R.J., Fairweather, P.G., 1995. Comparison of rapid methods for sampling the pipi, *Donax deltoides* (Bivalvia: Donacidae), on sandy ocean beaches. Marine and Freshwater Research 46: 1093–1099. https://doi.org/10.1071/MF9951093.

James, R.J., Fairweather, P.G., 1996. Spatial variation of intertidal macrofauna on a sandy ocean beach in Australia. Estuarine, Coastal and Shelf Science 43: 81–107. https://doi.org/10.1006/ecss.1996.0058.

Jaramillo, E., Pino, M., Filun, L., Gonzalez, M., 1994. Longshore distribution of *Mesodesma donacium* (Bivalvia: Mesodesmatidae) on a sandy beach of the South of Chile. The Veliger 37: 192–200.

McFadden, D., 1974. Conditional logit analysis of qualitative choice behaviour, in: Zarembka, P. (Ed.), Frontiers in Econometrics. Academic Press, New York, pp. 105–142.

McLachlan, A., Hanekom, N., 1979. Aspects of the biology, ecology and seasonal fluctuations in biochemical composition of Donax Serra in the East Cape. South African Journal of Zoology 14: 183–193. https://doi.org/10.1080/02541858.1979.11447668

McLachlan, A., Young, N., 1982. Effects of low temperature on the burrowing rates of four sandy beach molluscs. Journal of Experimental Marine Biology and Ecology 65: 275–284. https://doi.org/10.1016/0022-0981(82)90059-4.

McLachlan, A., Jaramillo, E., Defeo, O., Dugan, J., De Ruyck, A., Coetzee, P., 1995. Adaptations of bivalves to different beach types. Journal of Experimental Marine Biology and Ecology 187: 147–160. https://doi.org/10.1016/0022-0981(94)00176-E.

McLachlan, A., De Ruyck, A., Hacking, N., 1996. Community structure on sandy beaches: patterns of richness and zonation in relation to tide range and latitude. Revista Chilena de Historia Natural 69: 451–467.

Millar, R.B., Anderson, M.J., 2004. Remedies for pseudoreplication. Fisheries Research 70: 397–407. https://doi.org/10.1016/j.fishres.2004.08.016.

Murray-Jones, S., 1999. Conservation and management in variable environments: the surf clam, *Donax deltoides*. Unpublished PhD thesis., University of Wollongong, Wollongong.

Murray-Jones, S., Steffe, A.S., 2000. A comparison between the commercial and recreational fisheries of the surf clam, *Donax deltoides*. Fisheries Research 44: 219–233. https://doi.org/10.1016/S0165-7836(99)00095-8.

Murtaugh, P. A. (2007). Simplicity and complexity in ecological data analysis. Ecology 88: 56–62. https://doi.org/10.1890/0012-9658(2007)88[56:SACIED]2.0.CO;2.

Owner, D., Rohweder, D.A., 2003. Distribution and habitat of Pied Oystercatchers (*Haematopus longirostris*) inhabiting beaches in northern New South Wales. Emu 103: 163–170. https://doi.org/10.1071/MU01053.

R Core Team (2016). R: A Language and Environment for Statistical Computing. R Foundation for Statistical Computing, Vienna, Austria.

Schlacher, T.A., Schoeman, D.S., Dugan, J., Lastra, M., Jones, A., Scapini, F., McLachlan, A., 2008. Sandy beach ecosystems: key features, sampling issues, management challenges and climate change impacts. Marine Ecology 29 (Supplement 1), 70–90.

Short, A.D., 2007. Beaches of the New South Wales Coast, second ed. Sydney University Press, Sydney.

Soares, A.G., McLachlan, A., Schlacher, T.A., 1996. Disturbance effects of stranded kelp on populations of the sandy beach bivalve *Donax serra* (Röding). Journal of Experimental Marine Biology and Ecology 205: 165–186. https://doi.org/10.1016/S0022-0981(96)02595-6.

Totterman, S.L. 2019. Estimating the abundance of benthic macrofauna from across shore transects. Unpub. prepint. bioRxiv 542258. https://www.biorxiv.org/content/early/2019/02/06/542258

Venables, W.N., Ripley, B.D., 2002. Modern Applied Statistics with S, fourth ed. Springer, New York.

